# Molecular Logic of DNMT3A1 Recruitment: Resolving Multivalency at the Chromatin Interface

**DOI:** 10.64898/2026.04.27.721037

**Authors:** Drew McDonald, Emma Sachs, Talia Rapoport, Daniel Niizawa, Manxin Cao, Nataly Konechne, Elizabeth Lee, Norbert Reich

## Abstract

Histone modifications correlate with DNA methylation, but the underlying mechanisms remain unclear. The DNA methyltransferase DNMT3A1 engages nucleosomes through multivalent interactions with linker DNA, histone H3 tails, the acidic patch, and ubiquitinated H2A. Using binding assays, kinetic analyses, and nanopore sequencing of engineered nucleosomes, we show that DNMT3A1 binding reflects avidity, limiting the impact of individual contacts. Accordingly, disrupting single interactions minimally affects overall affinity, whereas isolated domains and truncations remain modification-sensitive. Although histone modifications have little effect on k_cat_/K_m_ for linker methylation, disrupting ADD-H3K4me0 interactions redistributes methylation away from the nucleosome core particle. Nucleosome competition assays reveal that H3K4me0 and H3K36me2 promote selective linker methylation, whereas H3K27me3 and PRC2/EZH2 have no effect. Notably, despite strong UDR-dependent binding to H2AK119ub1-modified nucleosomes, this mark fails to enhance methylation over an unmodified competitor. We propose a model of commitment to catalysis to reconcile weak kinetic differences with strong substrate selectivity. These findings highlight avidity and commitment in governing DNMT3A1 nucleosome recognition and DNA methylation specificity.

## Introduction

Epigenetic inheritance is governed by a tripartite architecture of DNA methylation, histone modification, and non-coding RNA (Skvortsova, and Bogdanović, 2018; Fitz-James and Cavalli, 2022). These pathways are known to converge to maintain genome stability and drive cellular identity (Putiri and Robertson, 2011; Wagner, 2025). The biophysical logic by which these distinct pathways communicate remains a frontier in molecular biology. Importantly, it has been shown that disruption of this crosstalk is a hallmark of some human diseases (Lempiäinen and Garcia, 2023). Yet the inherent complexity of these multi-component systems, defined by their combinatorial nature and reversibility, has long obscured the precise mechanisms of information transfer (Ruthenburg et al, 2007).

The *de novo* methyltransferase DNMT3A1 stands at the center of this complexity (Sendžikaitė et al, 2019; Kibe et al, 2021; Uehara et al, 2023; Chen et al, 2024). Unlike its paralog DNMT3B1, which manages global methylation during embryogenesis, DNMT3A1 is the primary architect of cell-type-specific gene expression (Manzo et al, 2017; Gu et al, 2018; Yagi et al, 2020). It functions as a multimeric scaffold, forming homo- and hetero-tetramers (often with the regulatory factors DNMT3L and DNMT3B3) that must integrate a deluge of chromatin signals (Jia et al, 2007; Guo et al, 2015; Zhang et al, 2018; Xu et al, 2020; Gutekunst et al, 2026). While structural studies have identified specific domains such as the ADD and PWWP domains as “readers” of histone tails, a fundamental question remains: How does DNMT3A1 prioritize competing or synergistic chromatin marks (post-translational modifications) to execute precise methylation patterns?

Current models are often contradictory. Some evidence suggests that histone-tail sensing (e.g., H3K4me0) primarily facilitates complex recruitment via outer monomers (DNMT3L) (Otani et al, 2009; Zhang et al, 2010), while other data indicate that these same marks allosterically regulate the catalytic activity of internal monomers (DNMT3A1) (Li et al, 2011; Guo et al, 2015; Yan et al, 2026). This dual role, scaffolding versus allostery, suggests a sophisticated “multivalent sensing” mechanism that has, until now, confounded traditional biochemical analysis (Ruthenburg et al, 2007). The multivalent nature of the DNMT3A1-nucleosome interface presents a significant investigative challenge. Beyond engaging linker DNA, DNMT3A1 achieves recruitment through a complex interplay with H3 histone tails, H2A ubiquitination, and the H2A/H2B acidic patch (Li, Chen and Lu, 2021; Bröhm et al, 2022; Chen et al, 2024; Gretarsson et al, 2024; Ward et al, 2026). How these individual contacts collectively dictate binding affinity and, more critically, DNA methylation specificity remains elusive. However, fundamental principles derived from other multivalent systems provide relevant insights, including (1) carbohydrate recognition by lectins (Roy et al, 2016; Guo et al, 2017; Martínez-Bailén et al, 2023), (2) cell membrane antigen recognition (Csizmar et al, 2019), and (3) polycomb repression (Ruthenburg et al, 2007; Justin et al, 2016; Davis et al, 2021; Gong et al, 2026). These studies suggest that such interactions are inherently dynamic. In these contexts, multivalency serves as a dual-purpose mechanism that simultaneously amplifies binding avidity and refines molecular discrimination.

In this study, we resolve these complexities by interrogating the interaction of DNMT3A1 with an array of modified nucleosomes featuring H3K4, H3K27, and H3K36 methyl-variants, alongside H2AK119 ubiquitination and acidic patch accessibility. Utilizing a high-resolution competition assay based on nanopore sequencing of 5-methylcytosine, we demonstrate that DNMT3A1 employs a hierarchical discrimination strategy. Our results reveal that H3K4me0 and H3K36me2 provide a sufficient and primary basis for nucleosome targeting and linker DNA methylation; however, H3K27me3 and H2AK119 ubiquitination fail to independently direct methylation in this context. We propose a unified framework where DNMT3A1 acts as a multivariate integrator, synthesizing signals from the DNA linker, the histone tails, and the nucleosome acidic patch to transform the histone code into permanent DNA methylation patterns (Ruthenburg et al, 2007).

## Results

### Multivalent interactions between DNMT3A1 and nucleosomal elements mask intrinsic histone modification specificity through avidity-driven binding

To answer how DNMT3A1 engages and discriminates between repressive and activating histone modifications, we bound full-length DNMT3A1 to different modified nucleosomes using AlphaLISA and anisotropy binding assay platforms. Full-length DNMT3A1 was used in order to preserve as much biologically relevant complexity as possible. In the AlphaLISA assay DNMT3A1 binds all modified and unmodified nucleosome substrates with K_d_ values ranging from 0.48 to 0.82 nM with minimal variation across substrates (SE ≤ 0.043 nM) (Fig. 1B). However, within this tight range of EC_50_ values, DNMT3A1 showed a binding preference hierarchy of H2AK119ub1 > H3K36me2 > unmodified > H3K27me3 > H3K4me3 which is consistent with published cellular studies (Noh et al, 2015; Manzo et al, 2017; Weinberg et al, 2019; Kibe et al, 2021; Yano et al, 2022; Chen et al, 2024). This narrow dynamic range of EC_50_ values suggests that the multivalent interactions of the DNMT3A1 on other nucleosome elements such as linker DNA and the acidic patch (Fig. 1A), mask the true binding discrimination to each histone modification. DNMT3A1 tetramer formation further amplifies interaction valency by enabling up to four subunits within a single complex to engage distinct nucleosomal features. Fluorescence anisotropy measurements of DNMT3A1 on FAM-tagged modified nucleosomes yielded similar binding trends with K_d_ values spanning from 3.68 to 6.56 nM (Fig. 1C). These results motivate the deconstruction of DNMT3A1 N-terminal regulatory domains to reveal the true binding energetics of DNMT3A1’s engagement with individual histone modifications.

**Figure 1.**
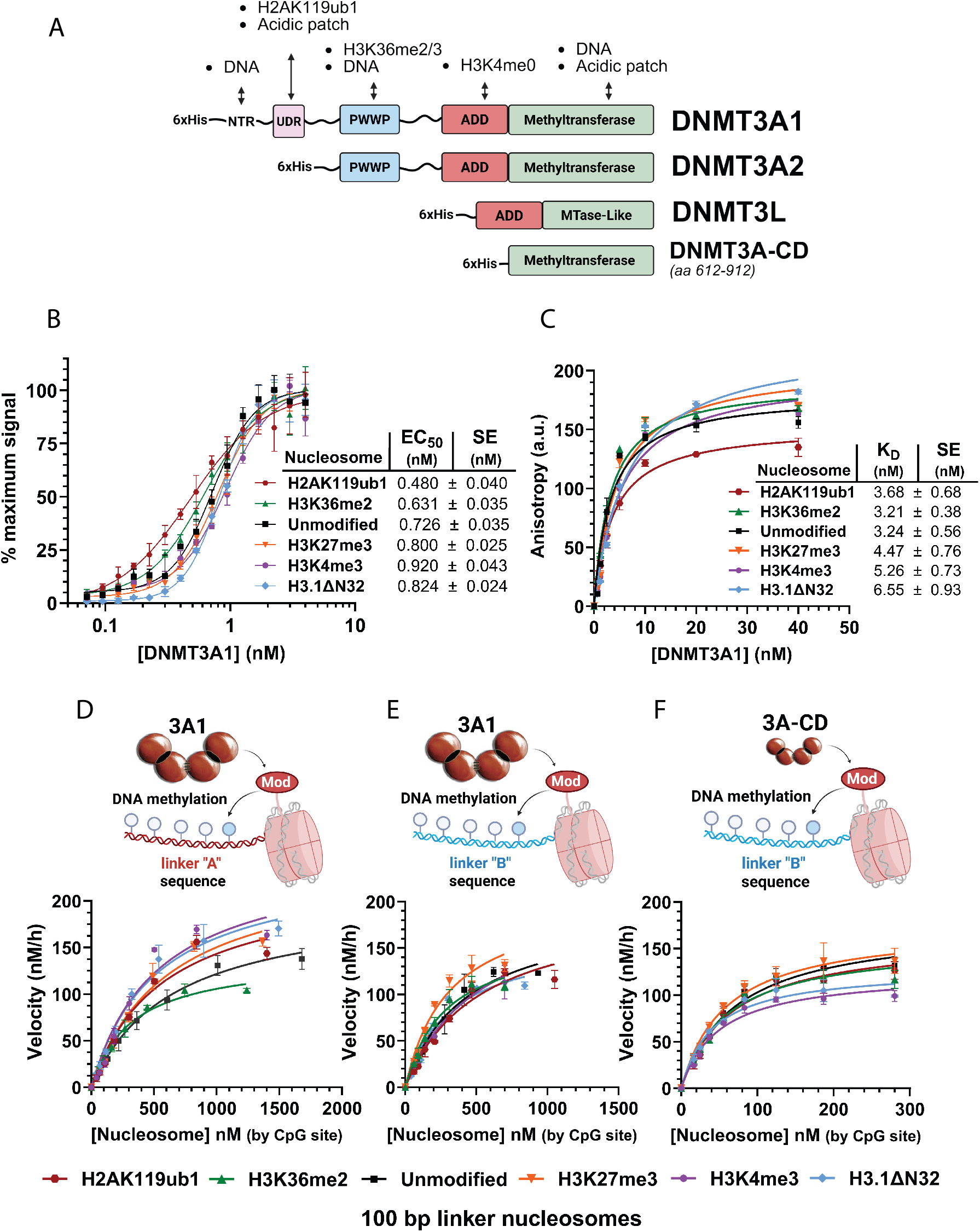
DNMT3A’s multivalent interactions with nucleosome elements veils DNMT3A’s binding preference and discrimination between histone modifications. (A) Domain map of the DNMT3A1 and its multivalent interactions with modified and unmodified nucleosome elements. DNMT3A2 splice isoform, DNMT3L homolog domain differences displayed along with a DNMT3A catalytic domain (DNMT3A-CD) protein construct used as a negative control in this study. (B) AlphaLISA binding assays of tetrameric DNMT3A1 on biotin-tagged modified 25 bp linker nucleosomes. Binding asymptotes and EC_50_ values were determined using four-parameter logistic curve fitting (R^2^ ≥ 0.97). Each curve represents data points preformed in triplicate; EC_50_ values are summarized to the right as mean ± SE. (C) Fluorescence anisotropy binding assay of tetrameric DNMT3A1 on FAM-tagged modified 25 bp linker nucleosomes. K_d_ values were determined using one-site binding curve fitting (R^2^ > 0.96). Measurements were performed in duplicate; K_d_ values are summarized at right as mean ± SE. (D) Michaelis-Menten plot of 50 nM DNMT3A1 on H2AK119ub1, H3K36me2, H3K27me3, H3K4me3, H3.1ΔN32 modified and unmodified 100 bp linker “A” nucleosomes quantified by CpG site. Kinetic parameters derived from Michaelis-Menten curve fitting (R^2^ ≥ 0.95) and are reported in Table 1 as mean ± SE. Each curve represents data collected in triplicate. (E) Same as D, using nucleosomes assembled with linker “B” DNA sequence. (F) Michaelis-Menten plot of 50 nM tetrameric DNMT3A catalytic domain on H2AK119ub1, H3K36me2, H3K27me3, H3K4me3, H3.1ΔN32, and unmodified 100 bp linker “B” nucleosomes quantified by CpG site. Kinetic parameters were obtained by Michaelis-Menten fitting (R^2^ ≥ 0.95) and are reported in Table 1 as mean ± SE.

### DNMT3A1 exhibits mildly enhanced catalytic efficiency on H3K36me2 modified nucleosomes but not on H2AK119ub1 modified nucleosomes

Next, we wanted to answer how histone modifications could regulate DNMT3A1 activity and catalytic efficiency on modified 100 bp linker nucleosomes (Fig. 1D,E and Table 1). To answer this, we conducted k_cat_/K_m_ analysis of DNMT3A1 and DNMT3A catalytic domain on modified nucleosomes with different linker substrates; linker “A” and linker “B”. These linkers were designed to enable a comparative analysis using nanopore sequencing (Table 1). All methylation assays were conducted in buffer conditions previously reported in (Ward et al, 2026) with NaCl (20 mM) for optimal enzyme activity (Holz-Schietinger and Reich, 2010; Holz-Schietinger et al, 2012). Across both linker DNA contexts (“A” and “B”), H3K36me2-modified nucleosomes consistently exhibited lower K_m_ apparent values relative to the unmodified nucleosomes as well as increased catalytic efficiencies (Fig. 1D,E and Table EV1). Based on k_cat_/K_m_ comparisons, DNMT3A1’s observed discrimination toward the H3K36me2-modified nucleosome is consistent with prior velocity data (Wapenaar et al, 2024). Wapenaar et al, 2024 report a more dramatic effect on velocity due to their use of high salt (150 mM NaCl) in the activity assay, which also resulted in near absence of activity on their free DNA substrates, at odds with prior studies showing good velocity with free DNA (Holz-Schietinger and Reich, 2010; Holz-Schietinger et al, 2012). When assessing DNMT3A’s catalytic efficiency (k_cat_/K_m_) on H2AK119ub1 modified nucleosomes, no significant enhancement relative to the unmodified control nucleosomes were observed (Fig. 1D,E and Table 1). Although the UDR domain in the DNMT3A1 has been demonstrated to contribute to enhanced binding to nucleosomes with the H2AK119ub1 modification (Fig. 4F) as well as previously reported (Chen et al, 2024; Wapenaar et al, 2024), this enhanced binding does not lead to enhanced K_m_ or catalytic efficiency. This finding suggests that UDR and Ub1 interactions do not allosterically activate DNMT3A1 on nucleosomes.

**Table 1:** Kinetic parameters of DNMT3A1 and the DNMT3A catalytic domain on modified nucleosomes and free DNA substrates. Kinetic parameters derived from Fig. 1D-F. DNMT3A-CD Velocity vs [Substrate] plot on linker “A” nucleosomes data not shown. All kinetic parameters derived in GraphPad Prism v10.6 using Michaelis-Menten curve fitting with R^2^ ≥ 0.95. Reactions were run with 50 nM tetrameric DNMT3A1 or DNMT3A-CD in 50 mM Tris (pH 7.8), 20 mM NaCl, 5 *μ*M [^3^H]-SAM (1:50 radiolabeled: unlabeled), 1 mM EDTA, 1 mM DTT, and 0.2 mg/mL BSA. DNMT3A1 + unmodified & H3.1ΔN32 linker “B”, and DNMT3A-CD + H2AK119ub1, H3K36me2, H3K27me3, and H3.1ΔN32 linker “A” nucleosomes curves run in duplicate. All other curves run in triplicate.

|  | Nucleosome Substrate | Nucleosome with 100bp Linker "A" DNA |  |  | Nucleosome with 100bp Linker "B" DNA |  |  |
| --- | --- | --- | --- | --- | --- | --- | --- |
| | | $k_{cat}$<br>($h^{-1}$) | $K_m$<br>($nM$)<br>(by CpG site) | Catalytic efficiency<br>$k_{cat}/K_m$<br>($h^{-1} \mu M^{-1}$) (by CpG site) | $k_{cat}$<br>($h^{-1}$) | $K_m$<br>($nM$)<br>(by CpG site) | Catalytic efficiency<br>$k_{cat}/K_m$<br>($h^{-1} \mu M^{-1}$) (by CpG site) |
| DNMT3A1 | Free DNA | 5.61 ± 0.36 | 480 ± 85 | 11.7 ± 2.2 | 4.67 ± 0.29 | 465 ± 82 | 10.0 ± 1.9 |
|  | Unmodified | 4.01 ± 0.24 | 647 ± 83 | 6.2 ± 0.9 | 3.89 ± 0.39 | 434 ± 90 | 9.0 ± 2.1 |
|  | H2AK119ub1 | 4.49 ± 0.35 | 568 ± 98 | 7.9 ± 1.5 | 3.97 ± 0.36 | 521 ± 91 | 7.6 ± 1.5 |
|  | H3K36me2 | 2.80 ± 0.11 | 323 ± 33 | 8.7 ± 0.9 | 3.32 ± 0.23 | 283 ± 43 | 11.7 ± 2.0 |
|  | H3K27me3 | 4.86 ± 0.32 | 617 ± 85 | 7.9 ± 1.2 | 4.15 ± 0.35 | 312 ± 54 | 13.3 ± 2.6 |
|  | H3K4me3 | 5.04 ± 0.43 | 529 ± 100 | 9.5 ± 2.0 | 4.24 ± 0.52 | 538 ± 120 | 7.9 ± 2.0 |
|  | H3.1ΔN32 | 4.85 ± 0.28 | 513 ± 65 | 9.5 ± 1.3 | 3.28 ± 0.33 | 317 ± 70 | 10.3 ± 2.5 |
| 3A Catalytic Domain | Free DNA | 3.21 ± 0.26 | 133 ± 23 | 24.1 ± 4.6 | 2.26 ± 0.12 | 38.6 ± 5.9 | 56.4 ± 7.9 |
|  | Unmodified | 2.56 ± 0.16 | 84 ± 12 | 30.5 ± 4.8 | 3.54 ± 0.15 | 71.7 ± 7.8 | 49.4 ± 5.8 |
|  | H2AK119ub1 | 3.93 ± 0.42 | 166 ± 32 | 23.7 ± 5.2 | 3.27 ± 0.16 | 65.9 ± 8.7 | 49.6 ± 7.0 |
|  | H3K36me2 | 3.84 ± 0.36 | 106 ± 21 | 36.2 ± 7.9 | 3.14 ± 0.16 | 58.8 ± 8.1 | 53.4 ± 7.8 |
|  | H3K27me3 | 3.04 ± 0.23 | 57.3 ± 11 | 53.1 ± 10.9 | 3.49 ± 0.15 | 58.7 ± 7.0 | 59.5 ± 7.5 |
|  | H3K4me3 | 2.45 ± 0.14 | 71.0 ± 10 | 34.5 ± 5.2 | 2.50 ± 0.11 | 49.9 ± 5.7 | 50.1 ± 6.1 |
|  | H3.1ΔN32 | 3.38 ± 0.32 | 84.7 ± 18 | 39.9 ± 9.3 | 2.58 ± 0.12 | 41.8 ± 5.8 | 61.7 ± 9.0 |

Both the transcriptionally activating histone mark H3K4me3 and the H3-tailless nucleosome control (H3.1ΔN32) resulted in reduced catalytic efficiencies (k_cat_/K_m_) of full-length DNMT3A1 on both linker “A” and linker “B” substrates (Fig. 1D,E; Table 1). Notably, an increase in k_cat_ was observed only for nucleosomes containing linker “A”, consistent with a redistribution of methylation away from the nucleosome core particle (NCP) (Fig. 3F). This effect is likely driven by the presence of more favorable CpG flanking sequences located 52 and 94 bp from the NCP in linker “A” (Table EV1). Together this demonstrates that DNMT3A1 activity is redirected away from the NCP when stabilizing histone tail interactions are disrupted.

H3K27me3-modified nucleosomes were observed to have a mild enhancement on catalytic efficiency when compared with unmodified nucleosomes for both DNMT3A1 and the DNMT3A1 catalytic domain (Fig. 1F and Table 1), suggesting activation is independent of DNMT3A1 regulatory domains. These data could suggest a potential indirect mechanism of H3K27me3 enhancing DNMT3A1 accessibility to DNA by altering H3 tail interactions with DNA and increasing linker DNA accessibility similar to mechanisms proposed previously (Bröhm et al, 2022; Ghoneim et al, 2021).

### Domain dissection of DNMT3A1 reveals binding discrimination of the PWWP domain to the H3K36me2 as well as the UDR domain to the H2AK119ub1 and acidic patch

To determine whether DNMT3A1 intrinsically discriminates H3K36me2-modified nucleosomes and to reconcile the lack of binding selectivity observed with full-length DNMT3A1 (Fig. 1B,C), we isolated the PWWP domain (DNMT3A1 aa 225-427) and assessed its binding to unmodified and H3K36me2-modified nucleosomes using AlphaLISA. Interestingly, isolating the PWWP domain showed a five-fold binding enhancement measured on the H3K36me2 modified nucleosomes compared to the unmodified nucleosome control (Fig. 2B). Thus, the removal of all other extraneous DNMT3A1 domains reduced the interaction multivalency, unmasking the significant contribution of the PWWP motif to H3K36me2 binding discrimination.

**Figure 2.**
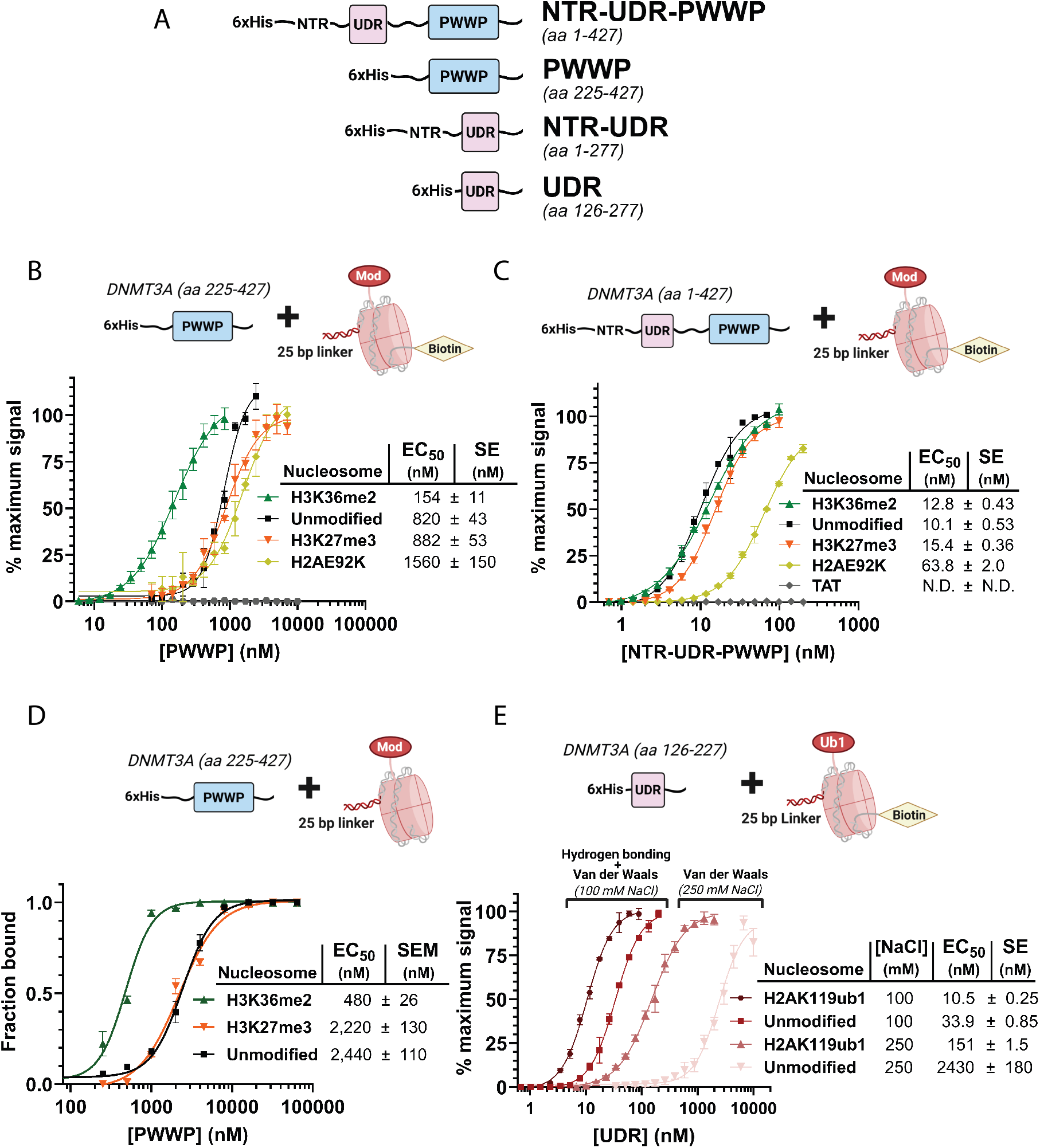
Domain analysis of DNMT3A1’s N-terminal regulatory domains reveal binding discrimination with H3K36me2 to PWWP domain, UDR to the acidic patch and H2AK119ub1, but no discrimination with H3K27me3 to the PWWP domain. (A) Domain map of DNMT3A1 C-terminal truncations constructs and isolated domains used in study. (B) AlphaLISA binding assay of PWWP domain (DNMT3A1 aa 225-427) to biotin-tagged H3K36me2, H3K27me3, H2AE92K modified and unmodified 25 bp linker nucleosomes and TAT as a negative control. Binding asymptotes and EC_50_ values were determined from four-parameter logistic curve fitting (R^2^ ≥ 0.97). Each curve represents data points preformed in triplicate; EC_50_ values are summarized to the right as mean ± SE. (C) AlphaLISA binding assay of NTR-UDR-PWWP domain (DNMT3A1 aa 1-427) to same constructs as in B. Binding asymptotes and EC_50_ values determined from four-parameter logistic curve fitting (R^2^ ≥ 0.98). Each curve represents data points preformed in triplicate; EC_50_ values are summarized to the right as mean ± SE. (D) Electrophoretic mobility shift assay (EMSA) quantification of (0.2 - 10 μM) PWWP domain (DNMT3A1 aa 225-427) on 10 nM of unmodified, H3K27me3, and H3K36me2 modified 25 bp linker nucleosomes. EMSA gels displayed in Fig. EV1. Fraction of nucleosome bound quantified from disappearance of unbound nucleosome bands. Mean ± SEM was collected from triplicate gels for each lane and fit with a four-parameter logistic curve fitting (R^2^ ≥ 0.95). EC_50_ values and SEM summarized to the right. (E) AlphaLISA binding assay of UDR wt (DNMT3A1 aa 126-277) on 2.5 nM of H2AK119ub1 modified and unmodified biotin-tagged 25 bp linker nucleosomes. Binding curves conducted in 100 mM to 250 mM NaCl to preserve and disrupt H-bonding energetics. Alpha signal asymptotes and EC_50_ values determined from four-parameter logistic curve fitting (R^2^ ≥ 0.97). Each curve represents data points preformed in triplicate; EC_50_ values are summarized to the right as mean ± SE.

We next assessed nucleosome binding using a C-terminally truncated DNMT3A1 (aa 1-427) construct containing the NTR, UDR and PWWP domains, which engage nucleosomes through the following multivalent interactions: (1) PWWP recognition of H3K36me2, (2) UDR contacts with the nucleosomal acidic patch, (3) and nonspecific DNA binding by the N-terminal Region (NTR) (Fig. EV2A-D). This protein construct bound all nucleosome substrates with a ten-fold enhanced affinity over the PWWP domain (DNMT3A1 aa 225-427) (Fig. 2B,C), demonstrating the UDR’s strong energetic contribution when interacting with the acidic patch of the nucleosome along with the NTR to DNA. Interestingly, the five-fold enhanced binding discrimination with the PWWP domain (DNMT3A1 aa 225-427) to the H3K36me2 nucleosome vanished either due to avidity, or a shift in the predominant interaction from the PWWP - H3K36me2 to the tighter UDR - acidic patch. This predominant UDR acidic patch interaction is also evident by the six-fold decrease in binding affinity of the NTR-UDR-PWWP to the acidic patch mutant H2AE92K nucleosome (Fig. 2C). The lack of H3K36me2 discrimination by DNMT3A1 (aa 1-427) may suggest that simultaneous engagement of the acidic patch (via the UDR) and H3K36me2 (via the PWWP domain) does not occur within a single DNMT3A1 monomer. Rather, these interactions could be partitioned across different subunits within the DNMT3A1 tetramer, with the distal subunit engaging the acidic patch and dimer pair recognizing the modified H3 tail, or vice versa.

To address the lack of significant discrimination for DNMT3A1 recognition of the H2AK119ub1 modified nucleosome substrate (Fig. 1B,C), the UDR (DNMT3A1 aa 126-277) was isolated from DNMT3A1 to minimize avidity and bound to H2AK119ub1 modified and unmodified biotinylated 25 bp linker nucleosomes using AlphaLISA (Fig. 2E). Binding assays were conducted in buffer conditions with high and low [NaCl] concentrations (250 mM and 100 mM) to reveal binding discrimination attributed primarily by van der Waals interactions between the UDR - H2AK119ub1 non-polar interface and the acidic patch (250 mM NaCl) as well as binding discrimination amidst hydrogen bonding, electrostatic, and van der Waals interactions between the UDR - H2AK119ub1 and acidic patch (Chen et al, 2024 Fig. S4). The UDR has an approximately three-fold enhancement in binding to the H2AK119ub1 modified nucleosome compared to the unmodified nucleosome control in 100 mM NaCl and an ever greater ~16-fold enhancement in 250 mM NaCl. This demonstrates that the binding discrimination between the UDR to the H2AK119ub1 modification can be hidden amidst the UDR’s hydrogen-bonding and electrostatic interactions with the nucleosome acidic patch. The interaction strength between the UDR and the unmodified nucleosome control diminished around ~70-fold from low to high NaCl conditions, while the UDR’s interaction to the H2AK119ub1 modified nucleosome diminished only around ~14-fold from the low to high NaCl conditions. These results demonstrate the salt-sensitivity of the hydrogen bond and ionic bond rich UDR - acidic patch interaction. The UDR - H2AK119ub1 interactions are persistent even amidst more stringent, higher NaCl conditions.

### The DNMT3A1 reader domain PWWP preferentially binds H3K36me2 but not H3K27me3 modified nucleosomes

To test whether or not the H3K27me3 modification can act as a scaffold for DNMT3A1 recruitment via binding in the PWWP domain, the PWWP domain (DNMT3A1 aa 225-427) was isolated and titrated against unmodified and H3K27me3 modified nucleosomes along with H3K36me2 modified nucleosome positive controls in two separate binding assays; AlphaLISA and EMSA (Fig. 2B,D). In both the AlphaLISA and EMSA, the PWWP binding to the H3K27me3 modified nucleosomes had an apparent affinity comparable to the unmodified nucleosomes, which suggests it likely does not serve a scaffolding role for recruiting the DNMT3A1 in the context of a nucleosome. In contrast, the PWWP domain exhibited a tighter affinity for H3K36me2 modified nucleosomes compared with unmodified nucleosome, consistent with previous studies (Fig. 2D; Fig. S1; Weinberg et al, 2019; Wapenaar et al, 2024).

### 5-methyl-C nanopore DNA sequencing is a viable platform for probing spatial distribution of nucleosome linker methylation by DNMT3A

To establish a platform for probing how histone modifications influence DNMT3A-mediated methylation across nucleosome linkers, methylation analysis was conducted on two unmodified control nucleosomes with 100 bp linkers (“A” and “B”) containing distinct flanking CpG sequences, as well as the corresponding free DNA (Table EV1). Differences in the preferences of CpG flanking sequences in linker “A” vs “B” by DNMT3A1 are most pronounced on free DNA (Fig. 3A; Handa and Jeltsch, 2005; Dossmann et al, 2024). The absence of optimal flanking sequences in linker “B” redirects methylation to more favorable CpG sites within the Widom 601 sequence (positions −24 and −70) on free DNA. Methylation analysis of linker “A” and “B” sequences when incorporated into unmodified nucleosomes show less flanking sequence preference and enriched methylation proximal to the NCP, regardless of CpG flanking sequence (Fig. 3C). Methylation on Widom 601 DNA wrapped around the nucleosome core is significantly suppressed which helps confirm proper Widom positioning on reconstituted nucleosomes. Dividing nucleosome linker methylation by free DNA methylation normalizes differences in methylation imparted by DNMT3A1 flanking sequence preference and accentuates effects of the nucleosome core particle (NCP) on DNMT3A1’s methylation across the DNA linker.

**Figure 3.**
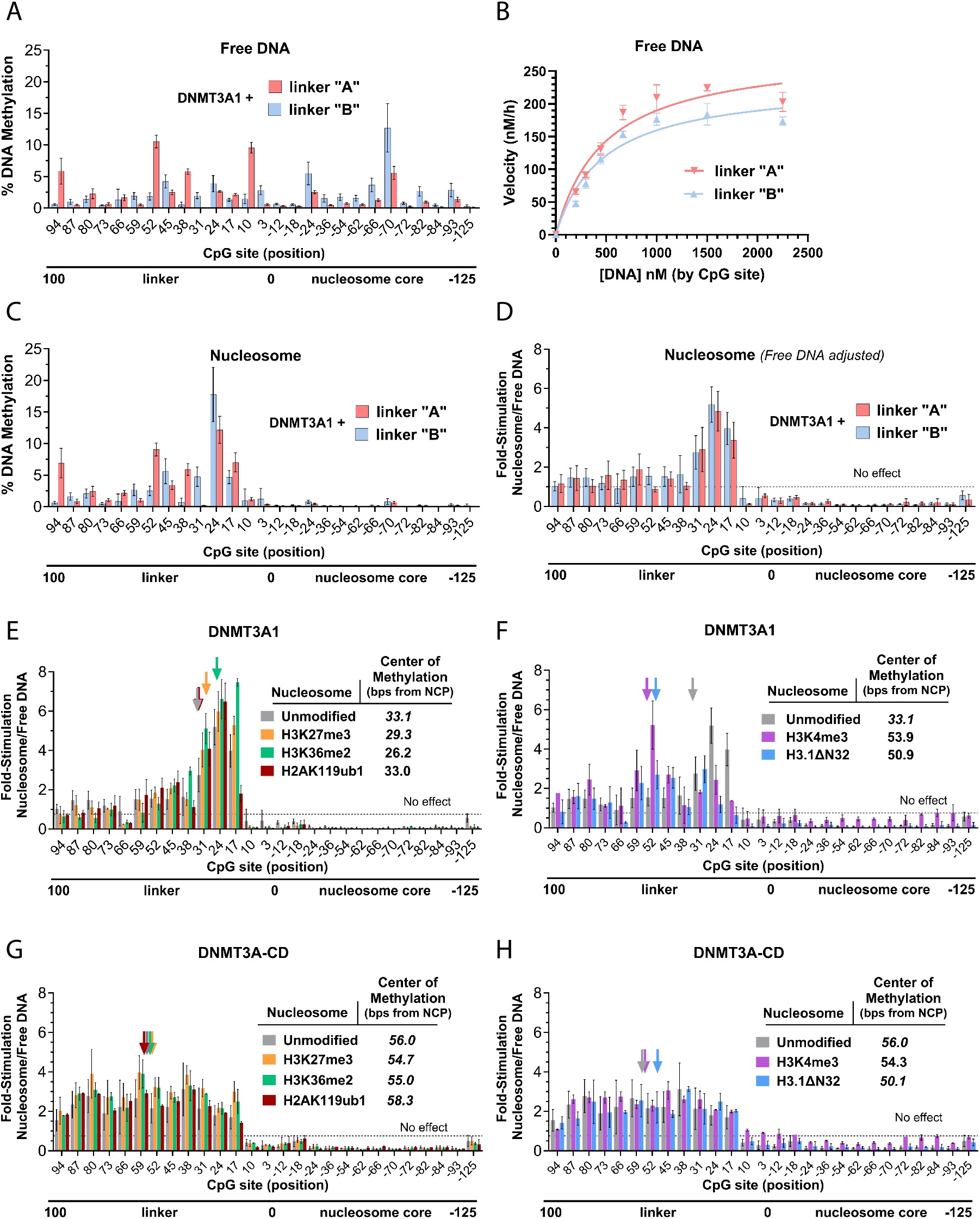
Site-specific DNA methylation of DNMT3A1 on modified nucleosomes show no change in linker methylation in the presence of H2AK119ub1 or H3K27me3 modifications. (A) DNMT3A’s site-specific 5-methyl-C DNA methylation on Widom 601-100 bp linker “free” DNA (unwrapped from histone octamer). Assay contained DNMT3A1 (125 nM tetrameric) with Widom 601-100 bp linker DNA (250 nM by DNA particle, 6750 nM by CpG site). Linker “A” and “B” contain same number of CpG sites and positions with different flanking sequences to the CpG sites. Methylation analysis determined using nanopore sequencing. (B) Michaelis-Menten plot of 50 nM DNMT3A1 tetramer on Widom 601-100 bp linker “A” and “B DNA. DNMT3A1 exhibits similar K_m_ values for both linker “A” and “B” K_m_ = 480 ± 85 nM (by CpG) linker “A”, K_m_ = 465 ± 82 nM (by CpG) linker “B”. Differences in K_cat_ (17%) attributed to preferential flanking sequence at position 94 and 52 and 10 on linker “A”. Curves run in technical triplicate. Additional kinetic parameters are displayed in Table 1 (C) Linker methylation of linker “A” and “B” by 125 nM DNMT3A1 tetramer on 100 bp linker “A” and linker “B” nucleosomes (250 nM by particle, 3500 nM by CpG site. CpG sites in Widom 601 excluded). Methylation analysis determined using nanopore sequencing. (D) Nucleosome % linker methylation by DNMT3A1 divided by % methylation of “free linker DNA (C / A). (E) Distribution of linker methylation by DNMT3A1 on H3K27me3, H3K36me2, H2AK119ub1 modified and unmodified 100 bp linker “B” nucleosomes. Presence of H3K36me2 modification shifts center-of-methylation 7 bp closer to the NCP compared to the unmodified control. H3K27me3 and H2AK119ub1 modifications did not significantly change the methylation patterns from the unmodified nucleosome control. (F) Same as (E) with H3K4me3 and H3 tailless (H3.1ΔN32) modified nucleosomes with unmodified control. (G) Same (E) with 125 nM of DNMT3A catalytic domain control. (H) Same as (F) with 125 nM of DNMT3A catalytic domain control. Experiments in (A) and (D-H) were performed in biological duplicate; values represent the mean ± S.D. All center-of-methylation (COM) values summarized to the right.

### H3K4me0 nucleosomes bearing H3K36me2, H2AK119ub1, H3K27me3 modifications all enrich linker methylation proximal to nucleosome core by DNMT3A1

To determine whether repressive histone modifications modulate DNMT3A1 positioning on nucleosomes, we examined linker methylation patterns on nucleosomes bearing H3K27me3, H3K36me2, or H2AK119ub1 modifications (Fig. 3E) (Weinberg et al, 2019; Weinberg et al, 2021). DNA methylation patterns revealed that H3K27me3 and H2AK119ub1 modified nucleosomes closely resembled unmodified nucleosomes, with maximal DNMT3A1 stimulation of methylation occurring proximal to the NCP. Center-of-methylation (COM) values (weighted methylation average across free DNA adjusted data sets) showed minimal shifts relative to unmodified controls. Due to the presence of the repressive H3K4me0 mark in addition to the H3K27me3, H2AK119ub1, and H3K36me2 modified nucleosomes, NCP enrichment effects may be hidden or confounded by DNMT3A1’s - H3K4me0 interaction in its ADD domain. However, despite the presence of this additional H3K4me0 repressive signal, H3K36me2-modified nucleosomes exhibited a mild but reproducible shift in the methylation toward the NCP, resulting in a COM approximately 7 bp closer to the NCP compared to the unmethylated nucleosome. This shift suggests H3K36me2 plays an important role in scaffolding DNMT3A1 to the H3K36me2 modified NCP for closely localized linker methylation. Assays were repeated with the DNMT3A catalytic domain to validate the specificity of these histone modification-dependent enrichments of methylation by DNMT3A1 (Fig. 3G). Recruitment effects mediated by H3 tail interactions, ubiquitin recognition and acidic patch binding via the UDR are lost in the absence of N-terminal regulatory domains (Fig. 3G,H). The DNMT3A catalytic domain (DNMT3A-CD) COM values on all modified nucleosome substrates were within 3 base pairs from unmodified control COM of 56 bp.

### Disruption of H3 tail interactions redistributes DNMT3A1 activity away from the nucleosome core

We next examined nucleosomes bearing the transcriptionally activating histone modification H3K4me3 that is known to disrupt the DNMT3A’s H3 tail interaction in the ADD domain. The H3 tailless (H3.1ΔN32) nucleosome control (Fig. 3F) and the H3K4me3 modified nucleosome showed significant changes in DNMT3A1-mediated methylation proximal to the NCP. Both H3K4me3 and H3.1ΔN32 nucleosomes were accompanied by increased relative methylation at distal linker positions. This redistribution resulted in an approximately 18 bp shift of the COM away from the nucleosome core relative to unmodified nucleosomes. This shift in methylation was similar to the magnitude of shift observed from the DNMT3A catalytic domain experiments (Fig. 3H). This result suggests that all H3K4me0 enrichment effects via DNMT3A1’s tail interaction are lost via disruption with H3K4 tri-methylation or through the removal of the H3 tail (H3.1ΔN32). Attenuation of H3 tail - ADD domain interactions on the nucleosome reveals that intact UDR - acidic patch interactions alone are insufficient to maintain enrichment of linker methylation proximal to the NCP. This suggests that the distal DNMT3A1 UDR interaction with the acidic patch may stabilize the binding of the DNMT3A1 to the nucleosome but does not affect the proximal methylation across the linker DNA.

Interestingly, DNMT3A-CD and DNMT3A1 exhibit similar COM at 50 and 51 bp from the NCP on H3 tailless (H3.1ΔN32) nucleosomes. All other modified or unmodified tailed nucleosomes consistently show a COM closer to 56 bp away from the NCP by DNMT3A-CD (Fig. 3F-H). This mild shift in methylation suggests H3 tails may be interacting with the linker DNA proximal to the NCP, as previously suggested (Ghoneim et al, 2021). This H3-tail linker DNA interaction could cause mild disruptions in DNMT3A-CD’s accessibility to the NCP proximal CpG’s due to the absence of N-terminal regulatory domain that can bind and release the H3 tail from the DNA.

### DNMT3A1 ADD, PWWP, and UDR mutants reveal H3 tail - dependent methylation near the nucleosome core

To corroborate the provocative result that the H3K4me3 modification is sufficient for disrupting DNMT3A’s enriched DNA methylation proximal to the NCP (Fig. 1F), DNMT3A1 mutants were generated to disrupt specific H3 tail and UDR interactions. We first measured binding of the DNMT3A2 splice isoform and D529A ADD domain mutant to unmodified (H3K4me0) nucleosomes using AlphaLISA (Fig. 4A). The D529A ADD domain mutant bound the unmodified nucleosomes with a ~five-fold reduction in affinity compared to the DNMT3A2 wt. This demonstrates a strong dependency of the H3 tail for DNMT3A’s engagement with nucleosomes.

**Figure 4.**
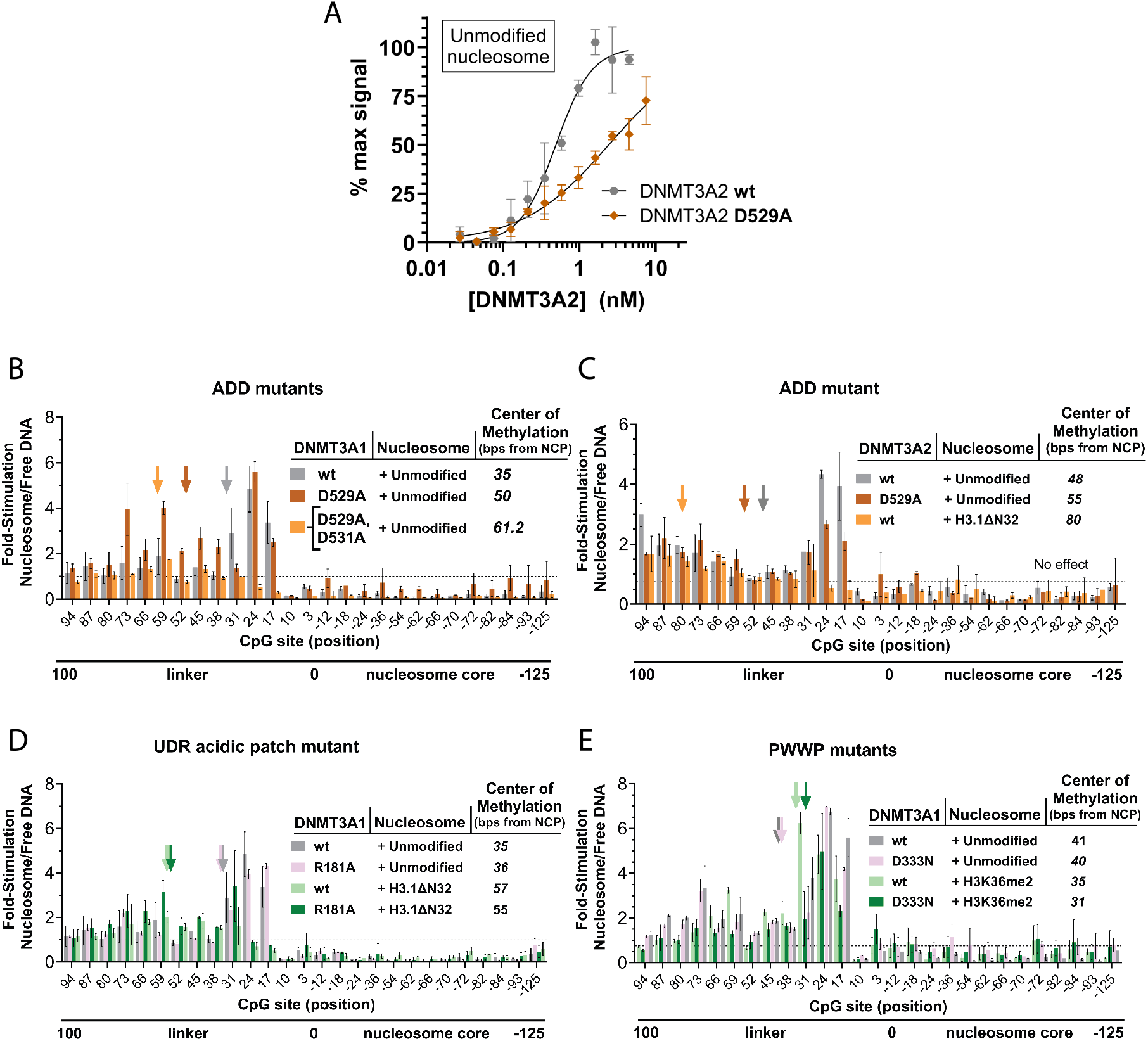
H3 tails are the predominant nucleosomal element responsible for enriching DNMT3A1 linker methylation proximal to the nucleosome core. (A) AlphaLISA binding assays of DNMT3A2 wt and the ADD domain mutant (D529A) on 2.5 nM biotinylated 25 bp linker unmodified nucleosomes. Binding asymptotes and EC_50_ values were determined by four-parameter logistic fitting (R^2^ ≥ 0.95). Each curve represents data collected in duplicate EC_50_ values are 0.49 ± 0.03 nM wt and 2.42 ± 0.18 nM (D529A) and are summarized at right as mean ± SE. (B) Distribution of linker methylation by tetrameric DNMT3A1 wt, and ADD domain mutants (D529A, D529A+D531A) on unmodified 100 bp linker “A” nucleosomes (C) Distribution of linker methylation by DNMT3A2 wt and ADD domain mutant (D529A) on unmodified 100 bp linker “A” nucleosomes as well as DNMT3A2 wt on H3 tailless (H3.1ΔN32) 100 bp linker “A” nucleosomes. (D) Distribution of linker methylation by DNMT3A1 wt, and UDR acidic patch mutant (R181A) on H3 tailless (H3.1ΔN32) and unmodified 100 bp linker “A” nucleosomes. (E) Distribution of linker methylation by DNMT3A1 wt, and PWWP domain mutant (D333N) on H3K36me2 modified and unmodified 100 bp linker “A” nucleosomes. All nanopore methylation assays were executed with 125 nM tetrameric DNMT3A1 or 3A2 enzyme on 100 bp linker “A” nucleosomes (250 nM by particle, 3500 by CpG site) and were performed in biological duplicate; values represent the mean ± S.D. Center-of-methylation (COM) values summarized to the right.

We next examined how disruption of the ADD domain affects the spatial distribution of linker methylation. On unmodified 100 bp linker nucleosomes, tetrameric DNMT3A1 wt displayed preferential methylation of CpG sites proximal to the NCP, reflected in a center-of-methylation (COM) value of 35 bp from the NCP (Fig. 4B). In contrast, the ADD mutants D529A and D529A + D531A exhibited progressive shifts in COM away from the NCP (COM = 50 and 61.2 bp, respectively), indicating loss of NCP-proximal enrichment despite an intact UDR domain. Thus, ADD-mediated engagement with the H3K4me0 tail is required for enriching methylation near the NCP via DNMT3A1, in the absence of other repressive histone modifications.

A similar requirement for the ADD domain was observed with DNMT3A2 (Fig. 4C). DNMT3A2 showed enrichment methylation at the NCP with a COM of 48 bp away from the NCP; whereas, the D529A mutant shifted linker methylation further away from the NCP (COM = 55 bp). Removal of the H3 tail (H3.1ΔN32) showed further shifting of the COM away from the NCP to 80 bp, which further demonstrates the dependency of the ADD H3 tail interaction for localizing the DNMT3A2’s methylation proximal to the NCP.

To determine whether UDR-dependent acidic patch binding drives NCP-proximal methylation, we compared linker methylation by the DNMT3A1 R181A mutant to wild type (Fig. 4D). Loss of acidic patch interaction did not alter NCP-proximal enrichment, as indicated by similar COM values (36 bp vs 35 bp). Even on H3-tailless nucleosomes, the R181A mutation did not shift linker methylation further from the NCP relative to wild type. Together with the distal shift observed for ADD domain mutants (Fig. 4B,C), these results indicate that H3K4me0 recognition by the ADD domain is the primary determinant of NCP-proximal methylation, whereas UDR-mediated acidic patch interactions alone are insufficient to drive this enrichment.

### Disruption of the PWWP domain does not change the enrichment of linker methylation proximal to H3K36me2 modified or unmodified nucleosome cores

We next examined whether disrupting the DNMT3A1 - H3K36me2 recognition by mutating the PWWP domain affects the DNMT3A1’s linker methylation positioning (Fig. 4D). The D333N PWWP mutant retained NCP-proximal enrichment on both unmodified and H3K36me2-modified nucleosomes, with COM values comparable to wild type (40 and 31 bp vs 41 and 35 bp, respectively). These results indicate that PWWP-dependent H3K36me2 recognition is not required for NCP-proximal methylation, likely because ADD-mediated H3K4me0 recognition is sufficient to direct positioning.

### Competition assay shows DNMT3A1 can discriminate between nucleosomes bearing the repressive H3K36me2 histone mark and the activating histone marks H3K4me3 but not H2AK119ub1 or H3K27me3 modified nucleosomes

To elucidate whether or not the DNMT3A1 activity can directly discriminate between nucleosomes bearing distinct histone modifications in the absence of additional cofactors, we used a nucleosome competition assay in which two otherwise identical nucleosome substrates differed only in linker DNA sequence and histone modification. Modified and unmodified nucleosomes were mixed at equal molar ratios in two-fold excess to the DNMT3A1 tetramer by particle. Nucleosomes were incubated with DNMT3A1 tetramer, after which site-specific DNA methylation on each linker was quantified by nanopore sequencing.

When modified linker “A” nucleosomes were competed against unmodified linker “B” nucleosomes, DNMT3A1 exhibited distinct methylation patterns depending on the histone modification (Fig. 5A,B). H3K36me2-modified nucleosomes showed a marked enrichment of DNA methylation across the linker relative to unmodified competitors, with particularly strong methylation at CpG sites proximal to the NCP. In contrast, H3K4me3-modified nucleosomes were consistently hypomethylated relative to unmodified nucleosomes across the linker, consistent with an inhibitory effect of H3K4 trimethylation on DNMT3A1 activity (Ooi et al, 2007; Otani et al, 2009; Zhang et al, 2010; Li et al, 2011). Nucleosomes bearing H3K27me3 or H2AK119ub1 exhibited linker methylation distributions that were largely comparable to unmodified nucleosomes, indicating little or no preferential targeting of methylation under these conditions. Similar trends were observed when the directionality of the assay was reversed, with unmodified linker “B” nucleosomes competed against modified linker “A” nucleosomes (Fig. 5C,D), confirming that linker sequence identity did not drive the observed preferences.

**Figure 5.**
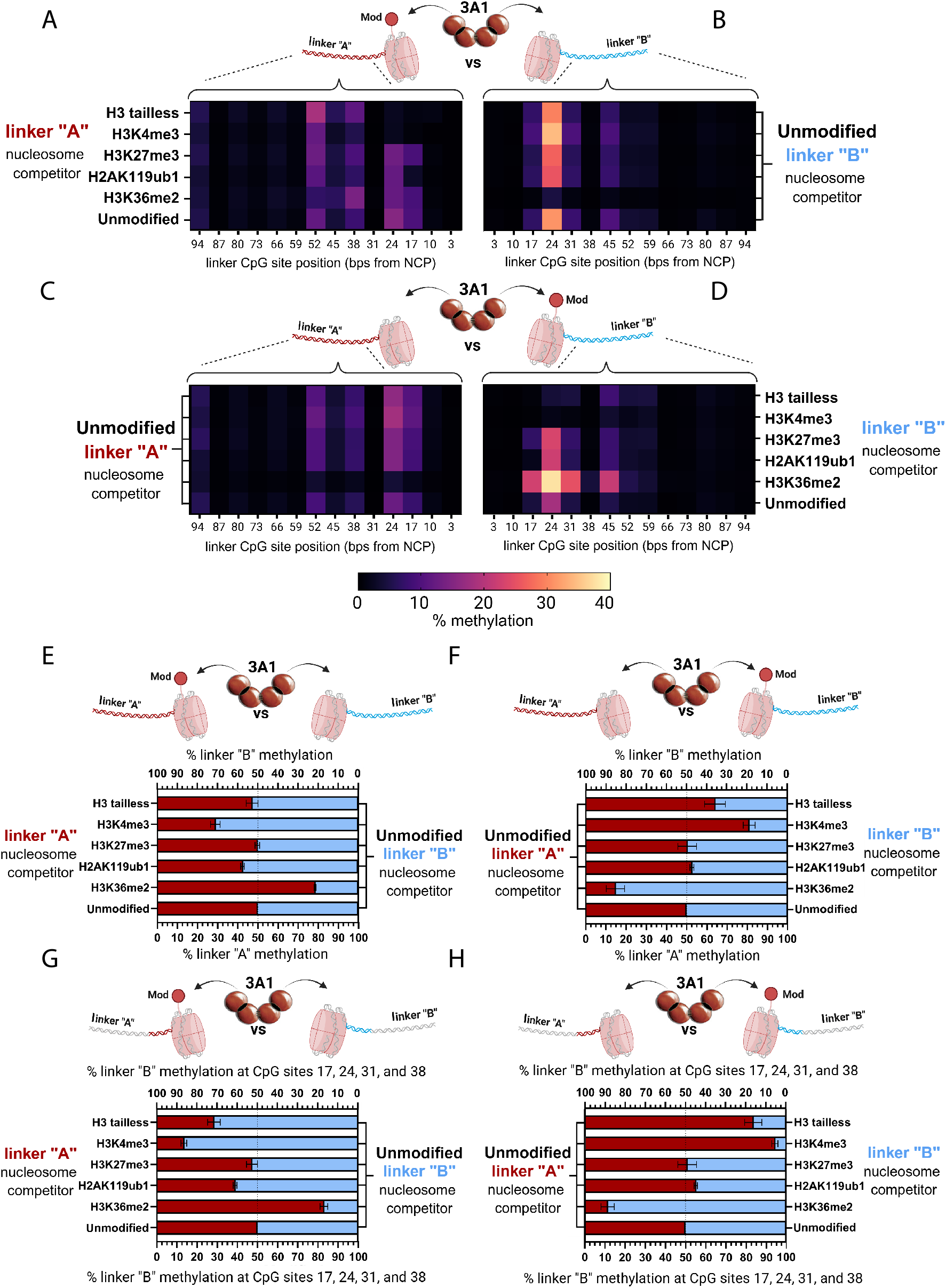
Nucleosome competition assay reveals preferential linker methylation by DNMT3A1 on H3K36me2 modified nucleosomes but not H2AK119ub1 or H3K27me3 modified nucleosomes. (A) Nucleosome competition assay site-specific linker methylation distributions measured on modified linker “A” nucleosomes competed against unmodified linker “B” nucleosomes. Assays contained 125 nM DNMT3A1 tetramer with 50/50 mixture of either H3.1ΔN32, H3K4me3, H3K27me3, H2AK119ub1, and H3K36me2 modified linker “A” nucleosomes paired with unmodified linker “B” nucleosomes (250 nM by particle, 3500 by CpG site). Methylation of linker “A” and linker “B” DNA mixture was determined via nanopore sequencing with separate alignments to the linker “A” and “B” sequences with thresholding of 0.8. (B) Site-specific linker methylation distribution measured on unmodified linker “B” nucleosomes competed against modified linker “A” nucleosomes. (C) Nucleosome competition assay site-specific linker methylation distributions measured on unmodified linker “A” nucleosomes competed against modified linker “B” nucleosomes. Same assay conditions used in A. (D) Site-specific linker methylation distributions measured on modified linker “B” nucleosomes competed against unmodified linker “A” nucleosomes. (E) Total methylation observed across (A) and (B) plotted in % methylation observed on modified nucleosomes with linker “A” vs unmodified nucleosomes with linker “B”. Nucleosome competition yielded 78.7 ± 0.12 % DNA methylation on H3K36me2 nucleosome, 42.2 ± 0.3 % on H2AK119ub1, 50.0 ± 0.8 % on H3K27me3, 29.2 ± 2.0 % on H3K4me3, and 47.4 ± 2.7 % on H3 tailless nucleosomes (F) Total methylation observed across (C) and (D) plotted in % methylation observed on modified nucleosomes with linker “B” vs unmodified nucleosomes with linker “A”. Nucleosome competition yielded 85.0 ± 4.3 % DNA methylation on H3K36me2 nucleosome, 47.3 ± 0.6 % on H2AK119ub1, 49.4 ± 4.3 % on H3K27me3, 18.9 ± 2.6 % on H3K4me3, and 35.6 ± 4.8 % on H3 tailless nucleosomes. (G) NCP proximal methylation observed at linker CpG sites 17, 24, 31, and 38 bp away from the NCP. Proximal methylation measured in (A) and (B) plotted in % methylation on modified nucleosomes with linker “A” vs unmodified nucleosomes with linker “B”. Nucleosome competition yielded 83.3 ± 1.5 % DNA methylation on H3K36me2 nucleosome, 39.2 ± 0.5 % on H2AK119ub1, 47.5 ± 2.5 % on H3K27me3, 13.7 ± 1.1 % on H3K4me3, and 28.7 ± 2.9 % on H3 tailless nucleosomes. (H) Same as G but with proximal methylation measured in (C) and (D) plotted in % methylation on modified nucleosomes with linker “B” vs unmodified nucleosomes with linker “A”. Nucleosome competition yielded 88.4 ± 3.0 % DNA methylation on H3K36me2 nucleosome, 44.5 ± 0.5 % on H2AK119ub1, 48.8 ± 4.3% on H3K27me3, 5.6 ± 1.2 % on H3K4me3, and 16.2 ± 4.0 % on H3 tailless nucleosomes. All experiments were performed in biological duplicate; values represent the mean ± S.D.

Quantification of total linker methylation across the competition assays revealed a definitive “preference” of the DNMT3A1 for methylating H3K36me2-modified nucleosomes over unmodified nucleosomes with 78.7 ± 0.12% of total methylation being depositing on H3K36me2 modified linker “A” nucleosomes, which increased to 85.0 ± 4.3% when modification was placed on linker “B” nucleosomes (Fig. 5E,F). In contrast, H3K4me3-modified nucleosomes captured only 29.2 ± 2.0% and 18.9 ± 2.6% of total methylation in the two assay orientations, respectively. H3K27me3 and H2AK119ub1-modified nucleosomes each accounted for approximately half of total methylation, similar to unmodified nucleosomes, indicating no strong preference by DNMT3A1. Analysis of methylation observed at CpG sites proximal to the NCP (17, 24, 31, and 38 bp away from the NCP) further accentuates the ability for H3K36me2 and the H3K4me0 histone modifications to enrich DNMT3A1’s methylation proximal to these modified repressive NCP loci.

Nucleosome competition assay results demonstrate that DNMT3A1 can directly and independently discriminate between nucleosomes bearing different histone modifications during DNA methylation. This is strong biochemical evidence that some histone modifications can directly influence the methylation activity of DNMT3A1 on linker DNA. H3K36me2 strongly promotes DNMT3A1-mediated linker methylation, particularly at nucleosome-proximal CpG sites, whereas H3K4me3 is inhibitory. In contrast, H3K27me3 and H2AK119ub1 do not confer preferential DNA methylation in this competition assay, indicating that these marks alone are not sufficient to bias DNMT3A1 activity in the absence of additional chromatin-associated factors.

### PRC2 forms a ternary complex with DNMT3A1 on nucleosomes but does not enhance DNMT3A1 catalytic activity or binding to nucleosomes

The relationship between H3K27me3 and transcriptional repression is well established in X-inactivation and stem cell differentiation (Tiwari et al, 2008; Singh et al, 2026). However, how H3K27me3 impacts DNA methylation, and the underlying mechanisms that alter DNMT3A1 activity, for example involving PRC2, EZH2 or H3K27me3, remain poorly investigated. To elucidate whether or not Polycomb repressive complex 2 (PRC2) modulates DNMT3A1 enzymatic activity on nucleosomes, we measured the V_max_ and K_m_ of the DNMT3A1 in the presence of PRC2 or its catalytic subunit EZH2 using unmodified 100 bp linker nucleosome substrates. Michaelis-Menten analysis revealed that neither PRC2 nor EZH2 had meaningful effects on the DNMT3A1’s catalytic efficiency on nucleosome substrates (Fig. 6A). In contrast, PRC2 increased the apparent K_m_ by roughly 2.5-fold, and 1.5-fold for the EZH2 on DNMT3A catalytic domain on nucleosomes, while having minimal effect on V_max_ (Fig. 6B). The elevated K_m_ observed for the DNMT3A catalytic domain likely reflects competitive interference with nucleosome or DNA binding. These effects were not observed with full-length DNMT3A1, suggesting that N-terminal regulatory domains mitigate PRC2-mediated interference with nucleosome engagement.

**Figure 6.**
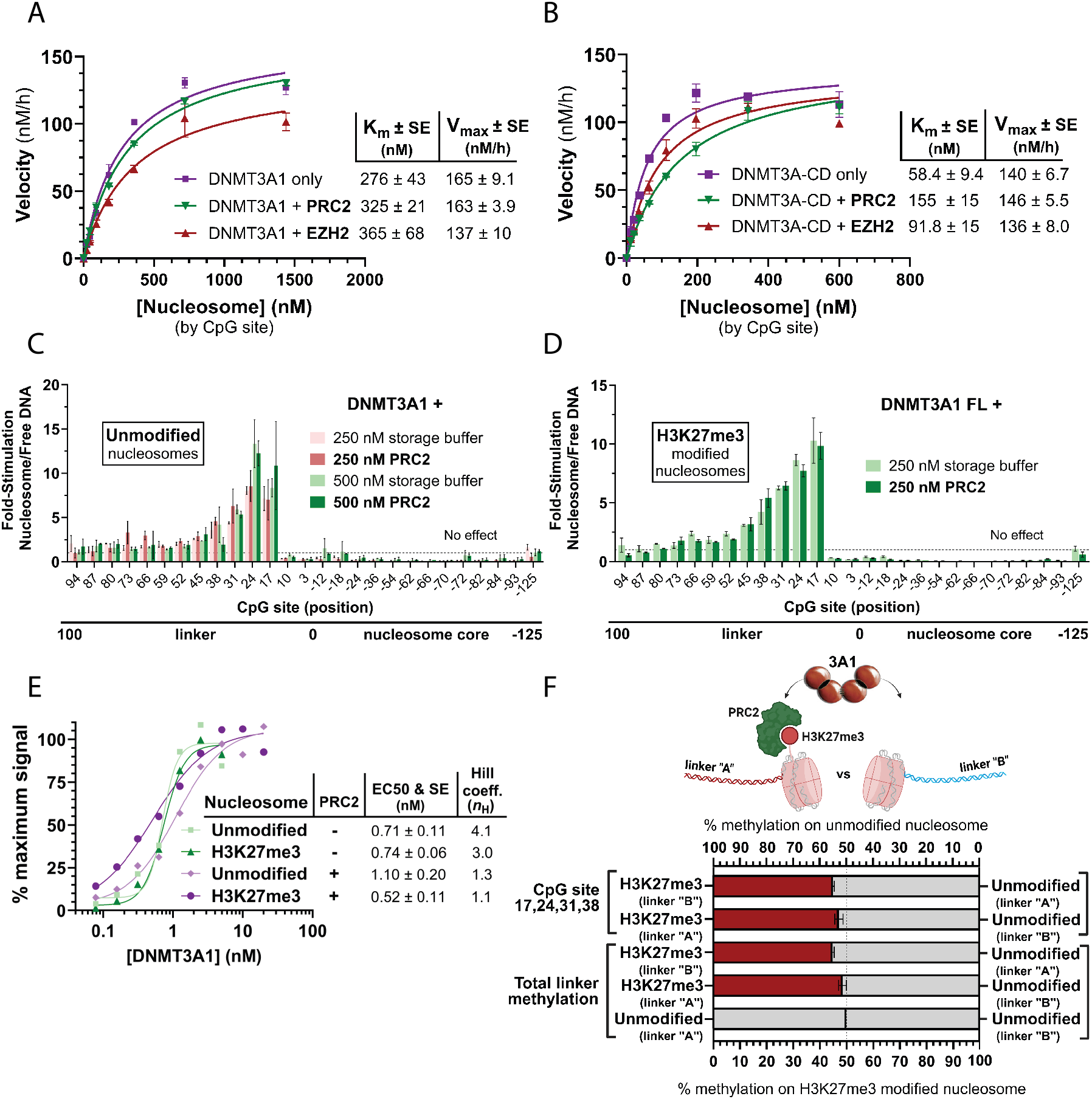
The ternary complex involving PRC2, DNMT3A and nucleosomes does not enhance DNMT3A1 binding or activity. (A) Michaelis-Menten plot of 50 nM DNMT3A1 tetramer on unmodified linker “A” nucleosomes with or without the presence of 200 nM PRC2 or EZH2. Kinetic parameters derived from Michaelis-Menten curve fitting (R^2^ ≥ 0.95). Kinetic parameters ± SE displayed to the right. Experiments run in technical duplicate (B) same as (A) with DNMT3A-CD catalytic domain negative control. (C) Distribution of linker methylation by 125 nM DNMT3A1 tetramer on unmodified 100 bp linker “B” nucleosomes (250 nM by particle, 3500 nM by CpG site) in the presence or absence of 250 nM or 500 nM of the PRC2. Methylation analysis determined using nanopore sequencing. Experiment was performed in biological duplicate; values represent the mean mean ± S.D. (D) Same as (C) with H3K27me3 modified nucleosomes with or without the presence of 250 nM PRC2 only. (E) AlphaLISA binding assay of DNMT3A1 on 2.5 nM of H3K27me3 modified or unmodified biotin-tagged 25 bp linker nucleosomes with or without the presence or absence of 200 nM PRC2. Binding curves conducted in AlphaLISA buffer conditions with 150 mM NaCl. Binding asymptotes and EC_50_ values determined from four-parameter logistic curve fitting (R^2^ ≥ 0.96). EC_50_ values, standard error (SE), and Hill Coefficients summarized to the right. (F) Nucleosome competition assay of 125 nM DNMT3A1 tetramer on 50/50 mixture of H3K27me3 modified and unmodified linker “A” and linker “B” nucleosomes pre-incubated with 250 nM PRC2. Assays contained 125 nM DNMT3A1 tetramer with 50/50 mixture of an individually modified linker “A and B” nucleosomes with unmodified linker “A and B” nucleosomes (250 nM by particle, 3500 nM by CpG site) preincubated with 200 nM PRC2. H3K27me3 modified nucleosome competitors showed 44.8 - 48.5 % DNA methylation against the unmodified nucleosome competitor. Experiment was performed in biological duplicate; values represent the mean ± S.D.

We next assessed whether PRC2 alters the spatial distribution of DNMT3A1-mediated DNA methylation along nucleosome linker DNA using nanopore sequencing. The addition of 250 nM or 500 nM PRC2 had no detectable effect on DNMT3A1’s site-specific linker methylation patterns on unmodified 100 bp linker nucleosomes (Fig. 6C). DNMT3A1 maintained a characteristic enrichment of methylation proximal to the NCP irrespective of PRC2 concentration. Similarly, PRC2’s engagement with H3K27me3 via the EED domain of the PRC2 did not alter the DNMT3A1’s magnitude or distribution of linker methylation on H3K27me3-modified nucleosomes (Fig. 6D), indicating that PRC2 binding does not redirect DNMT3A1 activity along linker DNA.

We then assessed whether or not the PRC2 acts as a scaffold for enhancing the DNMT3A’s engagement with nucleosomes at unmodified or H3K27me3 NCP loci. To answer this, we performed AlphaLISA binding assays with DNMT3A1 with unmodified or H3K27me3-modified nucleosomes in the presence and absence of PRC2, preincubated with the nucleosome substrates (Fig. 6E). Addition of PRC2 did not substantially weaken or enhance the DNMT3A1 binding affinity but resulted in a significant reduction in Hill-coefficient *n*_H_, decreasing from 4.1 to 1.3 on unmodified nucleosomes and from 3.0 to 1.1 on H3K27me3 nucleosomes. This reduction suggests a loss of cooperative DNMT3A1-nucleosome binding in the presence of PRC2, which is indicative of the formation of a ternary DNMT3A-PRC2-nucleosome complex.

Finally, we tested whether PRC2 influences the selectivity of nucleosome methylation in a direct nucleosome competition assay between H3K27me3 and unmodified nucleosome competitors. PRC2 was preincubated with a 50:50 mixture of H3K27me3-modified and unmodified nucleosomes. Leveraging PRC2’s enhanced binding affinity to H3K27me3 modified nucleosomes (Sanulli et al, 2015; Yu et al, 2019), the PRC2 - nucleosome complex formation should bias the H3K27me3 nucleosome with more PRC2 bound to the H3K27me3 modified nucleosome and less bound to the unmodified nucleosome. After the addition of the DNMT3A1, the total linker DNA methylation on each nucleosome competitor was measured using nanopore sequencing. PRC2 did not enhance DNMT3A1-mediated methylation of H3K27me3 modified nucleosomes relative to unmodified competitors, either at nucleosome proximal CpG sites or across the full linker region (Fig. 6F). These results indicate that PRC2 binding alone is insufficient to bias DNMT3A1 activity toward H3K27me3-marked nucleosomes.

Together, these data demonstrate that although PRC2 can co-occupy nucleosomes with DNMT3A1 and alter DNMT3A1 binding cooperativity, it does not alter DNMT3A catalytic activity, redistribute linker methylation, or confer preferential methylation of H3K27me3 modified nucleosomes under the experimental assay conditions tested in this study.

## Discussion

### Mechanistic insights into DNMT3A1 specificity and the histone code

The biochemical basis of DNMT3A1 specificity on nucleosomes has traditionally been viewed through the lens of recruitment, where histone modifications simply increase the local concentration of the enzyme via binding (Otani et al, 2009; Zhang et al, 2010; Ooi et al, 2007; Weinberg et al, 2021; Dukatz et al, 2019; Dhayalan et al, 2010). However, our results suggest a more nuanced kinetic model. While the “histone code” hypothesis posits that specific modifications drive DNA methylation patterns, many prior studies relied on isolated DNMT3A1 fragments or binding assays and did not comprehensively evaluate histone modifications effects on DNMT3A1’s catalytic turnover (Dhayalan et al, 2010; Dukatz et al, 2019; Weinberg et al, 2021; Chen et al, 2024; Gretarsson et al, 2024). By utilizing full-length DNMT3A1 under steady-state conditions with multiple catalytic cycles, we demonstrate that binding affinity (K_d_), catalytic efficiency (k_cat_/K_m_) and substrate discrimination do not always mirror one another.

Surprisingly, we observed only marginal changes in specificity constants for nucleosomes carrying well-characterized activating modifications, such as H3K4me0 or H3K36me2 (Fig. 1D-F and Table 1). This lack of dramatic discrimination in a non-competitive environment suggests that the “readout” of the histone code by DNMT3A1 is not a simple binary switch, but a sophisticated process governed by multivalent interactions (Davis et al, 2021; Ruthenburg et al, 2007).

### Multivalency and the “commitment” to catalysis

A central finding of this study is that full-length DNMT3A1 exhibits minimal binding discrimination between modified and unmodified nucleosomes, despite the known ability of its individual domains (e.g., PWWP, ADD) to recognize specific histone marks. We propose that this is a hallmark of multivalent avidity (Davis et al, 2021; Ruthenburg et al, 2007). Multivalent binding can result in dramatic affinity enhancements and additional specificity as well as retain dynamic features that allows for increased discrimination when compared to monovalent interactions (Mammen et al, 1998; Krishnamurthy et al, 2006; Davis et al, 2021). In a complex assembly like the DNMT3A1-nucleosome interactome, the total binding energy is summed across multiple interfaces including the DNA, the histone tails, the acidic patch, and ubiquitin moieties.

The results of our nucleosome competition assay (Fig. 5) suggest that this multivalency contributes to a “commitment” model of substrate discrimination (Johnson, 2010; Johansson et al, 2012; Banerjee et al, 2017; Scian et al, 2023). In enzymatic terms, commitment refers to the probability that an enzyme-substrate complex will proceed to catalysis rather than dissociate (Johnson, 2010; Johansson et al, 2012; Banerjee et al, 2017; Scian et al, 2023). Our competition assays reveal that while k_cat_/K_m_ values for individual nucleosomes with different histone modifications are similar, DNMT3A1 can effectively discriminate between some nucleosomes when they are presented simultaneously (Fig. 5). This competitive situation more realistically mimics the cellular environment (Xu et al, 2020). Thus, certain histone modifications (H3K4me and H3K36me) alter the partitioning of the complex, favoring release, followed by binding to more efficiently modified nucleosomes. Conversely, the lack of discrimination for H3K27me3 and H2AK119ub1 implies that these marks may either have a high degree of commitment, making release less likely, or require additional cellular factors or “readers” not present in our purified system to exert their biological effects.

### Decoding the impact of H3K27 and H2A ubiquitination

The relationship between H3K27 methylation and DNA methylation remains a subject of intense debate (Schlesinger et al, 2007; Brinkman et al, 2012; Reddington et al, 2013; Manzo et al, 2017; Weinberg et al, 2021). Our data provide a clear biochemical boundary: neither the presence of H3K27me3 nor the PRC2/EZH2 complex directly altered the catalytic activity or binding of DNMT3A1 in our assays (Fig. 6). This suggests that the correlations observed *in vivo* between H3K27 modifications and DNA methylation patterns (Li et al, 2021; Weinberg et al, 2021) likely stem from indirect recruitment mechanisms or chromatin remodeling events rather than a direct allosteric activation of DNMT3A1.

Furthermore, we must consider the “basal” state of our substrates. Most biochemical studies of histones, including ours, utilize nucleosomes in which H3K4me is unmethylated, which is well known to enhance DNMT3A1 binding (Otani et al, 2009; Zhang et al, 2010; Guo et al, 2015) (Fig. 4A; Fig. EV11) and linker methylation (Zhang et al, 2010). Given that H3K4me0 is itself a potent activator of DNMT3A1 via the ADD domain, it is possible that this “ground state” activation masks the subtler effects of other modifications which will require additional studies.

### Sequence-specific preferences and linker methylation

By integrating nanopore sequencing, we bridged the gap between ensemble kinetics and site-specific activity. The distinct methylation patterns observed at positions 52 and 94 in linker “A” (Fig. 3A) align with previous reports of flanking sequence preferences for the DNMT3A catalytic domain (Handa and Jeltsch, 2005; Dossmann *et al*, 2024). This indicates that while histone modifications provide the “scaffold” for recruitment and commitment, the fine-scale distribution of DNA methylation is ultimately refined by the local DNA sequence context. This dual layer of control involving epigenetic “priming” via histone tails and sequence-level “execution,” ensures the high fidelity of the DNA methylome.

## Materials and Methods

### Expression of the catalytic domain (DNMT3A-CD)

hDNMT3A-CD was expressed in NiCo21-(DE3) cells (NEB) using a codon-optimized plasmid pET28a-hDNMT3A-CD (Δ1-611). Cell cultures were grown in Luria Broth medium at 37°C to an OD_600_ = 0.8. Cultures were cooled to 25°C, then expression was induced by the addition of 1 mM IPTG (Gold Biotechnology) and cultured at 220 RPM. Cells were harvested by centrifugation after 5 h, and stored at −80°C.

### Expression of full-length DNMT3A1/A2 wt and point mutants

hDNMT3A1 was expressed in NiCo21-(DE3) cells (NEB) using a pET28a-hDNMT3A1 plasmid. Cell cultures were grown in Luria Broth medium at 37°C to an OD_600_ = 0.7. Cultures were cooled to 18°C, induced by the addition of 0.4 mM IPTG (Gold Biotechnology) and cultured at 220 RPM. Cells were harvested by centrifugation after 16 hours and stored at −80°C.

### Expression of DNMT3L

hDNMT3L was expressed in NiCo21-(DE3) cells (NEB) using a pTBY1-hDNMT3L plasmid with a stop codon at the end of the 3L gene. Cell cultures were grown in Luria Broth medium at 37°C to an OD_600_ = 0.6. Cultures were cooled to 28°C, induced by the addition of 1 mM IPTG (Gold Biotechnology) and cultured at 220 RPM. Cells were harvested by centrifugation after 4 h, and stored at −80°C.

### Expression of DNMT3A1 C-terminal truncations and isolated domains

hDNMT3A1 aa 1-277, aa 225-427, aa 1-427 were expressed in NiCo21-(DE3) cells (NEB) using pET28a-hDNMT3A1 aa 1-277, pET28a-hDNMT3A1 aa 225-427, and pET28a-hDNMT3A1 aa 1-427 plasmids. Cell cultures were grown in Luria Broth medium with 0.4% glucose at 37°C to an OD_600_ = 0.7. Cultures were cooled to 18°C, induced by the addition of 0.4 mM IPTG (Gold Biotechnology) and cultured at 220 RPM. Cells were harvested by centrifugation after 18 hours and stored at −80°C.

### Expression of histone octamers

The polycistronic coexpression and nondenaturing purification of histone octamers were performed as described previously (35). Briefly, pET29a-YS14 (Addgene) was transformed into Rosetta (DE3) pLysS (Novagen) cells and cultures were grown to OD_600_ = 0.4 at 37°C. The cultures were induced with 0.4 mM IPTG, grown for 20 hours at 220 RPM, harvested via centrifugation, and stored at −80°C.

### Expression of human EZH2 using baculovirus

Recombinant baculovirus was generated using the Bac-to-Bac baculovirus expression system (Thermo Fisher) with pFastBac/Flag-tev-EZH2 and pFastBac/ EZH2-tev-Flag transfer vectors. Flag-tagged EZH2 was expressed by infecting 150 mL ExpiSf9 insect cells (1 × 10^6^ cells/ml) in ExpiSf CD Medium (Thermo Fisher) with 300 μL P1 baculoviral suspension a (1:500), followed by a 70 h incubation. Cells were then harvested via centrifugation at 500 x g, and stored at −80°C.

### Purification of DNMT3A1 catalytic domain (DNMT3A-CD)

DNMT3A catalytic domain (Δ1-611) was purified as described previously (Ward et al, 2026). Clarified lysates were loaded into a 5 mL HisTrap excel column (Cytiva), washed extensively, and eluted with high imidazole. Eluted protein was concentrated and buffer-exchanged into 50 mM Tris (pH 7.8), 200 mM NaCl, 1 mM DTT, 1 mM EDTA, and 20% (v/v) glycerol. Protein concentration was determined by UV absorbance using ε_280_= 39,670 M^−1^cm^−1^ (ProtParam), assuming a monomeric subunit. Purified protein was flash-frozen in liquid nitrogen and stored at −80°C.

### Purification of full-length DNMT3A1 and 3A2 wt and point mutants

Full-length DNMT3A1 or 3A2 (wild-type and point mutants) was purified as described previously (Ward et al, 2026). Clarified lysates were loaded into 5 mL HisTrap excel column (Cytiva) with stepwise imidazole washes followed by gradient elution. Fractions containing DNMT3A1 were pooled, concentrated, and further purified by size-exclusion chromatography (Superdex 200, Cytiva) equilibrated in 50 mM Tris (pH 7.8), 200 mM NaCl, 1 mM DTT, 1 mM EDTA, and 10% (v/v) glycerol. Peak fractions were pooled and concentrated. Protein concentration was determined by UV absorbance using ε_280_= 142,010 M^−1^cm^−1^ (ProtParam), assuming a monomeric subunit. Purified protein was flash-frozen in liquid nitrogen and stored at −80°C.

### Purification of DNMT3L

NiCo21-(DE3) cells expressed with the pTBY1-hDNMT3L plasmid (modified with a stop codon at the end of the 3L gene) were resuspended in lysis buffer containing 50 mM Tris (pH 7.8), 500 mM NaCl, 25 mM imidazole, 0.2% Triton X-100, 1 mM PMSF, 25 μg/mL DNase I, and 10% (v/v) glycerol. The cells were lysed by sonication and the supernatant was clarified via centrifugation three times sequentially at 18,000 x g for 20 minutes. Lysates were applied to an ÄKTA start FPLC system (GE Healthcare) and loaded onto a 1 mL HisTrap excel column (Cytiva). The column was washed with 35 mL of 50 mM Tris (pH 7.8), 500 mM NaCl, 10% (v/v) glycerol and 25 mM imidazole. The protein was eluted with a 15 mL gradient of 25 mM – 500 mM imidazole in a 50 mM Tris (pH 7.8), 500 mM NaCl, 10% (v/v) glycerol. Eluted fractions were analyzed with SDS-PAGE and those with the highest purity were pooled, concentrated, and buffer swapped with a 10 kDa MW Amicon filter (Sigma) into 50 mM Tris (pH 7.8), 200 mM NaCl, 1 mM DTT, 1 mM EDTA, 20% (v/v) glycerol. The concentration of DNMT3L was determined using UV-vis with ε_280_ = 68,380 M^−1^ cm^−1^ derived by ProtParam. Concentrated protein was then flash frozen in liquid nitrogen for storage at −80°C.

### Purification of PWWP domain, UDR domain, and DNMT3A1 C-terminal truncations

NiCo21-(DE3) cells expressed with using pET28a-hDNMT3A1 aa 225-427, pET28a-hDNMT3A1 aa 126-277, pET28a-hDNMT3A1 aa 1-427 or pET28a-hDNMT3A1 aa 1-227 plasmids were resuspended in lysis buffer containing 25 mM NaH_2_PO_4_ (pH 7.8), 400 mM NaCl, 15 mM imidazole, 2 mM β-me, 0.1% Triton X-100, 1 mM AEBSF, 1 × Pierce EDTA-free PI Tablet (Thermo Fisher), 25 μg/mL DNase I, and 10% (v/v) glycerol. The cells were lysed with sonication and the supernatant was clarified via centrifugation three times sequentially at 18,000 x g for 20 minutes. Lysates were applied to an ÄKTA start FPLC system (GE Healthcare) and loaded onto a 5 mL HisTrap HP column (Cytiva). The column was washed with 50 mL of 25 mM NaH_2_PO_4_ (pH 7.8), 500 mM NaCl, 15 mM imidazole, 2 mM β-me, and 10% (v/v) glycerol. The protein was eluted with a 30 mL gradient of 15 mM – 300 mM imidazole in 50 mL of 25 mM NaH_2_PO_4_ (pH 7.8), 500 mM NaCl, 15 mM imidazole, 2 mM β-me, and 10% (v/v) glycerol. Eluted fractions were analyzed with SDS-PAGE and those with the highest purity were pooled and concentrated using a 10 kDa MW Amicon filter (Sigma). The protein was then applied to a 16/60 HiLoad column (GE Pharmacia) packed with Superdex 200 (Cytiva) equilibrated in 50 mM Tris (pH 7.8), 200 mM NaCl, 1 mM DTT, 1 mM EDTA, 10% (v/v) glycerol on an ÄKTA purifier FPLC system (GE Healthcare). Protein fractions were analyzed with SDS-PAGE for purity. Fractions with the highest purity were pooled and concentrated using a 10 kDa MW Amicon filter (Sigma). The concentration of DNMT3A1 C-terminally truncated proteins was determined using UV-vis with ε_280_ = 69,690 M^−1^ cm^−1^ for DNMT3A1 (aa 1-427), 19,605 M^−1^ cm^−1^ for DNMT3A1 (aa 1-277), and 50,085 M^−1^ cm^−1^ for DNMT3A1 (aa 225-427), derived by ProtParam. Concentrated protein was then flash frozen in liquid nitrogen for storage at −80°C.

### Purification of histone octamers

Histone octamers were purified from Rosetta (DE3) pLysS cells as described previously (Shim et al, 2012; Ward et al, 2026). Cells expressing histones were lysed in 50 mM Tris (pH 8.0), 2 M NaCl, 1 mM TCEP, 1 mM PMSF, and 30 mM imidazole, sonicated, and clarified by sequential centrifugation. Lysates were applied to a HisTrap column, washed, and eluted with a 30–500 mM imidazole gradient. Fractions containing all four histones were pooled, concentrated, and cleaved with thrombin to remove tags. Samples were further purified on a Superdex 200 column in 20 mM Tris (pH 8.0), 2 M NaCl, and 5 mM BME, pooled, supplemented with 10% glycerol, and flash-frozen for storage at −80°C. The concentration of histone octamer was determined using UV-vis with an ε_280_ = 44,700 M^−1^ cm^−1^ extinction coefficient.

### Purification of Flag-tagged EZH2

0.4 L of ExpiSf9 cells infected with recombinant Flag-tagged containing baculovirus resuspended in lysis buffer containing 50 mM Tris (pH 7.8), 500 mM NaCl, 0.3% Triton X-100, 1 mM PMSF, 1× Pierce Protease Inhibitor Tablet (Thermo Fisher), and 5% (v/v) glycerol. The cells were lysed with sonication and the supernatant was clarified via centrifugation at 13,000 × g for 30 minutes. 0.5 mL of Anti-Flag M2 Affinity Gel (Sigma-Aldrich) was equilibrated with lysis buffer and added to clarified lysate and rocked gently for 2 h. Beads were sedimented via centrifugation at 500 x g and were washed 4 times with 10 mL of 50 mM Tris (pH 7.8), 500 mM NaCl, 0.3% Triton X-100, and 5% (v/v) glycerol. They were washed an additional 4 times with 1 mL of the same buffer. Protein was eluted from beads 7 × times, incubating for 15 min with 200 uL of 250 ng/µL 3× FLAG peptide (Sigma-Aldrich) supplemented in 50 mM Tris (pH 7.8), 500 mM NaCl, 0.3% Triton X-100, and 5% (v/v) glycerol. Elutions were pooled and applied to a 16/60 HiLoad column (GE Pharmacia) packed with Superdex 200 (Cytiva) equilibrated in 50 mM Tris (pH 7.8), 200 mM NaCl, 1 mM DTT, 1 mM EDTA, 10% (v/v) glycerol on an ÄKTA purifier FPLC system (GE Healthcare). Protein fractions were analyzed with SDS-PAGE for purity. Fractions with the highest purity were pooled and concentrated using a 10 kDa MW Amicon filter (Sigma). The concentration of Flag-tagged EZH2 protein was determined using UV-vis with ε_280_ = 77,875 M^−1^ cm^−1^ derived by ProtParam. Concentrated protein was then flash frozen in liquid nitrogen for storage at −80°C.

### Preparation of nucleosome DNA

Nucleosome DNA constructs were generated as described previously (Ward et al, 2026). Briefly, a MluI site was inserted into the Widom 601 sequence in pGEM-3z/601 (Addgene). Tail PCR amplification with Q5 High-Fidelity polymerase was used to sequentially construct the Widom 601-100 bp linker “A” and linker “B” DNA template amplicons. These templates were used to generate 601-100, FAM-601-25, and biotin-601-25 molecules, which were purified and quantified with UV-vis using extinction coefficients of ε_260_ = 3.16 μM^−1^cm^−1^ for 601-100 DNA, 2.20 μM^−1^cm^−1^ for biotin-601-25 bp DNA, 2.27 μM^−1^cm^−1^ for FAM-601-25 DNA.

### Reconstitution of nucleosomes

Nucleosomes were reconstituted with histone octamers as described previously (Dyer et al, 2003; Paintsil and Morrison, 2023; Ward et al, 2026). Briefly DNA and octamers were mixed at 1:1 - 1:3 molar ratios in 20 mM Tris (pH 7.5), 2 M NaCl, 1 mM EDTA, and 1 mM DTT, incubated at 4°C, and then desalted over 40 - 48 h using a linear NaCl gradient. Reconstituted nucleosomes were incubated at 45°C to position octamers, concentrated using 100 kDa Amicon filters and supplemented with 10% glycerol. Concentrations were determined by UV-vis after separating DNA from histones. Integrity and purity were confirmed by SDS-PAGE, EMSA, and MluI digestion. Positioning on linker nucleosomes was further validated using a DNMT3A1 methylation nanopore assay (Ward et al, 2026).

### AlphaLISA binding assay

Biotin tagged nucleosomes or DNA substrates were prepared at a concentration of 10 nM in binding buffer consisting of 20 mM Tris (pH 7.8), (100 mM NaCl for DNMT3A1 domain isolates or 150 mM NaCl for full-length 3A1 or 3A2), 0.01% BSA, 0.01% NP-40 and 0.5 mM DTT. His-tagged interacting proteins were prepared in same buffer conditions at varying concentrations. 5 μL of the biotinylated substrate as well as 5 μL of the interacting protein were mixed together in a 384-well Alpha plate (Revvity) and incubated for 30 minutes at room temperature. Then 10 μL of 2.5 μg/mL Ni-Chelate AlphaLISA Acceptor beads (Revvity) and 5 ug/mL Streptavidin AlphaScreen Donor beads (Revvity) in 20 mM HEPES (pH 7.8), (100 mM NaCl for DNMT3A1 domain isolates or 150 mM NaCl for full-length 3A1 or 3A2), 0.01% BSA and 0.01% NP-40 was added to each well. The plate was incubated at room temperature in darkness for 1 hour. The AlphaLISA counts were measured using a TECAN SPARK equipped with Alpha Technology module (680 nm/750 mW laser excitation, 622.5 nm emission filter ± 25 nm bandwidth). All reactions were performed in triplicate. Curves and asymptotes were determined using Prism v10.6.1 Sigmoidal, 4PL, X is concentration curve fitting. Hook effect data points were excluded in the curve fitting when necessary.

### Fluorescence anisotropy binding assay for determination of K_d_

Changes in fluorescence anisotropy from binding interactions were measured using a TECAN SPARK microplate reader equipped with excitation and emission polarizers. Assays were performed in a 96-well plate containing 50 mM Tris (pH 7.8), 1 mM EDTA, 1 mM DTT, 0.2 mg/mL BSA, 20 mM NaCl, 10 *μ*M SAH (Sigma), 5 nM FAM-tagged substrate and varied amounts of enzyme in 50 *μ*L reactions. Measurements were taken after a 15-minute incubation at 25°C using an excitation of 485 nm and an emission of 520 nm. Data was background subtracted and fit to a One site - Specific binding model using Prism v10.6.1 unless otherwise stated. The K_d_ was determined using a 95% confidence interval.

### Radiochemical methylation assay

Methylation assays were used to determine nanomolar amounts of methylated DNA product. Reactions were carried out in 50 mM Tris (pH 7.8), 1 mM EDTA, 1 mM DTT, 0.2 mg/mL BSA, 20 mM NaCl, and 5 *μ*M SAM (Sigma Aldrich). The SAM cofactor was diluted with [^3^H]-SAM (PerkinElmer) in 10 mM H_2_SO_4_. Reactions were activated by the addition of nucleosome to a final volume of 20 *μ*L and incubated at 37°C for 30 minutes before they were quenched with 10 *μ*L of 0.3% SDS. Aliquots of reactions were spotted on Hybond-N+ membranes (GE Healthcare), then washed with 50 mM K_2_HPO_4_/KH_2_PO_4_ at pH 7.8 and ethanol. Membranes were dried and counted using a Hidex LS 300 scintillation counter. The k_cat_ was determined by dividing the velocity in nM/h by the enzyme concentration. The Michaelis-Menten curves were modeled using Prism v10.6.1 where V_max_ and K_m_ were extracted using a 95% confidence interval.

### Nanopore sequencing methylation assay and bioinformatic analysis

Methylation assays for nanopore 5mC sequencing contained 50 mM Tris (pH 7.8), 1 mM EDTA, 1 mM DTT, 0.2 mg/mL BSA, 20 mM NaCl, and 20 *μ*M SAM; with varied concentrations of MTase, free DNA or nucleosome. Reactions were activated with the addition of DNA or nucleosome and incubated at 37°C for 30 minutes. Reactions were quenched with 0.15% SDS and DNA was purified using a PureLink PCR Purification kit. Samples were sent to the UC Berkeley DNA Sequencing facility for library preparation, sequencing, base-calling and alignment to reference sequences. Libraries were prepared using the Rapid Barcoding Kit (SQK-RPD114-96, Oxford Nanopore Technologies) and loaded on a FLO-MIN114 flow cell (Oxford Nanopore Technologies) and sequenced using a MinION Mk1D (Oxford Nanopore Technologies). Raw sequencing reads were base-called for 5mC with Dorado v11.7.2 (Oxford Nanopore Technologies) and Remora, then aligned to target sequences using minimap2 (Li, 2018). Sequences were concatenated using Samtools (Li et al, 2009; Danecek et al, 2021) and pileup was done using Modkit (Oxford Nanopore Technologies, 2025) with a 0.8 threshold filter for cytosine and 5mC. When applicable, center-of-methylation (COM) values derived from weighted methylation average across free DNA adjusted data sets. Fold stimulation values < 1 were excluded from the analysis.

### Nucleosome competition assay

Two separate batches of both modified and unmodified histone octamers were wrapped with two separate 601-100 bp linker “A” and linker “B” nucleosome substrates as a form of DNA barcoding. Modified linker “A” nucleosomes and unmodified linker “B” nucleosomes were mixed at 50:50 molar ratios at 125 nM each for a total of 250 nM of nucleosomes per solution. Modified linker “B” mixtures with unmodified linker “A” nucleosomes were also formed at 50:50 molar ratios as well. Nanopore methylation assay solutions comprised of 125 nM of DNMT3A1 in 50 mM Tris (pH 7.8), 1 mM EDTA, 1 mM DTT, 0.2 mg/mL BSA, 20 mM NaCl (20, 50, 80 mM for Fig. EV4), and 20 *μ*M SAM were aliquoted and reactions initiated with the addition of the 50:50 nucleosome mixture and incubated at 37°C for 30 minutes. Reactions were quenched with 0.15% SDS and DNA was purified using a PureLink PCR Purification kit. Competition assay DNA samples were sent to the UC Berkeley DNA Sequencing Facility for library preparation, sequencing, base-calling and alignment to reference sequences. Nanopore sequencing procedures were followed above, however alignments were done with both the 601-100 bp linker “A” template sequence and the 601-100 bp linker “B” for each sample mixture to assess the methylation that occurred on the modified vs unmodified nucleosome.

### Electrophoretic mobility shift assay (EMSA)

For EMSA assays with nucleosomes, 10 nM of 25 base pair nucleosomes and varying concentrations of PWWP protein were prepared in 50 mM Tris (pH 7.8), 0.05 mg/mL BSA, 75 mM NaCl, 1 mM DTT, 1 mM EDTA, 0.05% (v/v) Triton X-100, 20% (v/v) glycerol, and 0.01% Bromophenol blue. Samples were preincubated together for 30 min at 4 °C and separated via 5% 29:1 acrylamide Tris native-PAGE gels with 0.5× TBE for 2 hours at 4 °C. Gels were stained with 1:5000 SYBR gold (ThermoFisher) for 20 min and imaged with UV excitation using a Gel Doc EZ imager (Bio-Rad).

For free DNA EMSA binding assays, 40 nM of 601-25 bp DNA was incubated with varying concentration of DNMT3A1 C-terminally truncated protein at RT for 20 min in 50 mM Tris (pH 7.8), 1 mM EDTA, 1 mM DTT, 20 mM NaCl, 0.05% Tween-20, and 5% (v/v) glycerol. Reactions were diluted with 2× Native Sample Buffer (Bio-Rad) and separated via 6% 75:1 acrylamide Tris native-PAGE gels with 0.5× TBE for 90 min at 4 °C. Gels were stained with 1:5000 SYBR gold (ThermoFisher) for 20 min and imaged with UV excitation using a Gel Doc EZ imager (Bio-Rad).

### Purchased modified octamers, nucleosomes and PRC2 complex

The following recombinant histone octamers were purchased from Epicypher and used in nucleosome reconstitutions for this study: Unmodified (ca.16-0001), H3.1ΔN32 tailless (ca.16-8016), H3K27me3 (ca.16-8317), H3K4me3 (ca.16-8316), H3K36me2 (ca.16-8319), and H2AK119ub1 (ca.16-8395). Purchased nucleosomes from Epicypher include biotinylated H2AE92K linker nucleosomes (ca. 16-1030). Recombinant PRC2 complex (EZH2, SUZ12, EED, RbAp46, and RbAp48) was purchased from Active Motif (ca. 31387)

## Data Availability

All expression vector sequences used in this study are accessible at: https://doi.org/10.5281/zenodo.19394786

Uncropped gel images accessible at: https://doi.org/10.5281/zenodo.19396628

Raw alphaLISA and anisotropy data accessible at: https://doi.org/10.5281/zenodo.19490394 Raw kinetic data accessible at: https://doi.org/10.5281/zenodo.19490025

Nanopore sequencing and competition assay data accessible at: https://doi.org/10.5281/zenodo.19672014

## Author Contributions

**Drew McDonald:** Conceptualization; Project administration; Methodology; Investigation; Data curation; Formal analysis; Writing - original draft and editing. **Emma Sachs:** Methodology; Investigation; Data. **Talia Rapoport:** Methodology; Investigation; Data. **Daniel Niizawa:** Methodology; Investigation; Data. **Manxin Cao:** Investigation; Data curation. **Nataly Konechne:** Investigation; Data curation. **Elizabeth Lee:** Methodology. **Norbert Reich:** Conceptualization; Supervision; Funding acquisition; Writing - review and editing.

## Acknowledgments

We acknowledge Ethan Ward and Raymond Abdo for their methodology. We also thank the UC Berkeley DNA Sequencing Facility for library preparation and nanopore 5mC sequencing. Research was enabled by the National Science Foundation under Grant No. 2403840.

